# A lentiviral toolkit to monitor airway epithelial cell differentiation using bioluminescence

**DOI:** 10.1101/2024.02.09.579635

**Authors:** Jessica C. Orr, Asma Laali, Pascal F. Durrenberger, Kyren A. Lazarus, Marie-Belle El Mdawar, Sam M. Janes, Robert E. Hynds

**Affiliations:** Lungs for Living Research Centre, UCL Respiratory, University College London, London, U.K; Epithelial Cell Biology in ENT Research (EpiCENTR) Group, Developmental Biology and Cancer Department, Great Ormond Street UCL Institute of Child Health, University College London, London, U.K; UCL Cancer Institute, University College London, London, U.K

**Keywords:** airway epithelium, lentiviral transduction, basal cells, ciliated cells, mucosecretory cells, primary cell culture

## Abstract

Basal cells are adult stem cells in the airway epithelium and regenerate differentiated cell populations, including the mucosecretory and ciliated cells that enact mucociliary clearance. Human airway basal cells can proliferate and produce differentiated epithelium *in vitro*. However, studies of airway epithelial differentiation mostly rely on immunohistochemical or immunofluorescence-based staining approaches, meaning that a quantitative approach is lacking. Here, we use a lentiviral reporter gene approach to transduce primary human basal cells with bioluminescence-based reporter constructs to monitor airway epithelial differentiation longitudinally. We generated three constructs driven by promoter sequences from the *TP63*, *MUC5AC* and *FOXJ1* genes to quantitatively assess basal cell, ciliated cell and mucosecretory cell abundance, respectively. We validated these constructs by tracking differentiation of basal cells in air-liquid interface and organoid (‘bronchosphere’) cultures. Transduced cells also responded appropriately to stimulation with interleukin 13 (IL-13; to increase mucosecretory differentiation and mucus production) and IL-6 (to increase ciliated cell differentiation). We anticipate that these constructs will be a valuable resource for researchers monitoring airway epithelial cell differentiation in primary epithelial and/or induced pluripotent stem cell (iPSC) cell cultures.

**Graphical Abstract:** 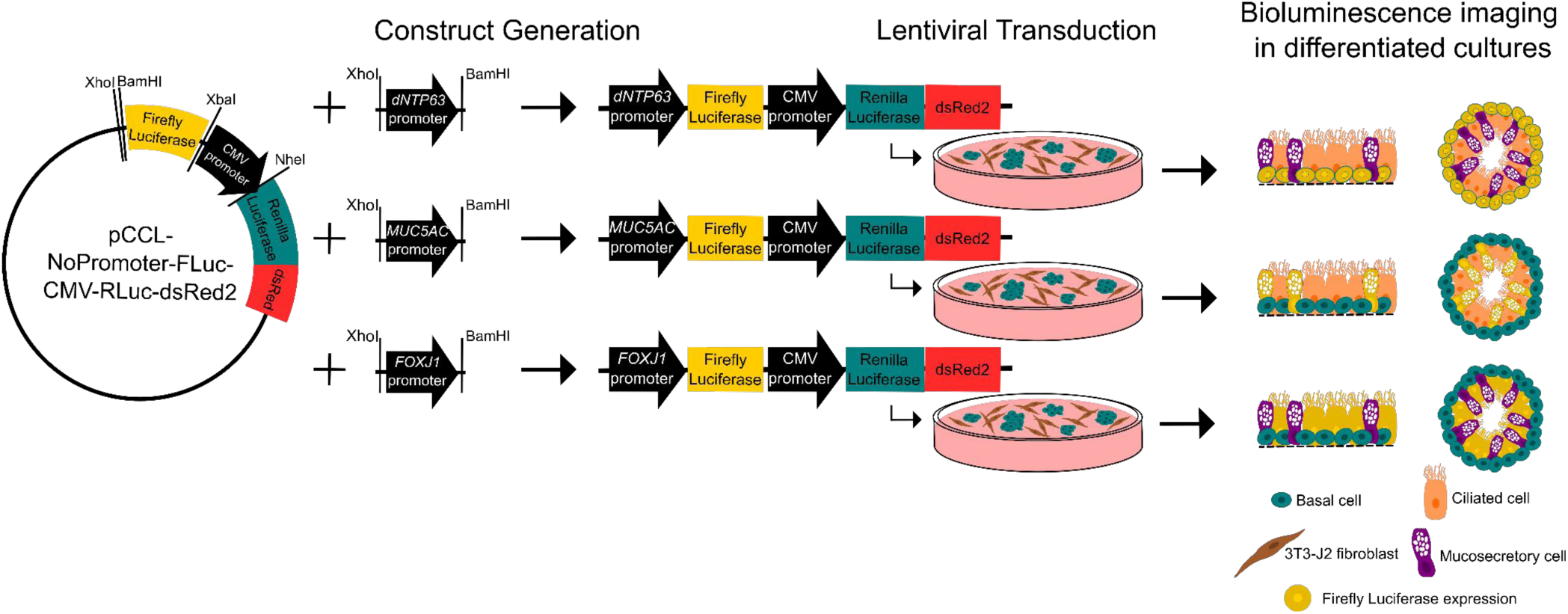

## INTRODUCTION

Airway epithelial cells play a critical role in maintaining respiratory function by forming a protective barrier against inhaled pathogens, toxins, and environmental insults. Basal stem cells, mucosecretory cells, and ciliated cells are the most abundant cell types of the airway epithelium (1). Basal cells serve as adult stem cells and play a crucial role in maintaining tissue homeostasis by repairing the damaged epithelium (2). Mucosecretory cells and ciliated cells each enact aspects of the mucociliary escalator, which acts to trap inhaled particles and remove them from the respiratory tract. Mucosecretory cells produce and secrete airway mucus, while ciliated cells produce motile force to move the mucus layer proximally and remove trapped matter from the respiratory system (3).

Understanding the normal development, homeostasis and repair of the airway epithelium is crucial to understand how these processes are altered in disease states, to develop new respiratory medicines and in the pursuit of airway tissue engineering approaches. Primary airway epithelial cell culture can achieve long-term propagation of patient-derived cell cultures and the *in vitro* differentiation of airway basal cells to differentiated cell types (4, 5). However, studies of airway epithelial cell differentiation typically rely on either immunohistochemical and immunofluorescence staining, which are limited in their ability to provide dynamic and quantitative data, or indirect measures, such as monitoring transepithelial electrical resistance (6). The introduction of reporter genes, for example fluorescent proteins or luciferase enzymes, can enable quantitative readout of biological activity. This is commonly achieved by transfection of plasmids, but reporter cell lines can be created by transduction with lentiviral reporter constructs. Lentiviruses transduce proliferating and non-proliferating cell types and integrate the construct into the genome for stable expression. Bioluminescence imaging captures light produced by the oxidation of substrates by luciferase enzymes. The most commonly used are firefly luciferase (from the North American firefly, Photinus pyralis), which uses D-luciferin as its substrate and releases yellow-green light, and renilla luciferase (from the sea pansy, Renilla reniformis), whose substrate is coelenterazine and releases predominantly blue light (7).

Tumor protein p63 (TP63) is a transcription factor that is expressed by basal cells in stratified squamous epithelia and the pseudostratified airway epithelium. Transactivating and N-terminally truncated (ΔNp63) TP63 isoforms are generated from two promoter sequences, generating at least 11 distinct isoforms (8). Airway basal cells predominantly express the ΔNp63α isoform (9). Consistent with its role in maintaining epithelial stem cells in other tissues, knockdown of *TP63* inhibits proliferation and differentiation, and promotes basal cell senescence (10). Expression of *TP63* is restricted to a subset of undifferentiated airway basal cells and reduces upon commitment to differentiation (11–14). The mucin 5AC (*MUC5AC)* gene encodes a secreted, polymeric mucin, and transcripts are restricted to airway goblet cells in the human tracheal and bronchial epithelium (15). Tobacco smoking induces *MUC5AC* expression and *MUC5AC* expression is elevated in the airways of patients with chronic obstructive pulmonary disease (COPD) (16). Variants in *MUC5AC* have been associated with chronic sputum production (17), asthma (18) and pulmonary fibrosis (19). Forkhead box protein J1 (FOXJ1) is a transcription factor involved in the late stages of ciliogenesis and, in the airways, is expressed uniquely by multiciliated cells (20). As a result of the cell type specificity of *TP63*, *MUC5AC* and *FOXJ1* expression, we reasoned that these would represent suitable promoter sequences to drive luciferase reporter gene expression in lentiviral reporter constructs to quantify the abundance of basal cells, mucosecretory, and multiciliated cells, respectively.

## MATERIALS AND METHODS

### Luciferase gene reporter constructs

To generate an editable promoter-reporter construct vector (pCLL-NoPromoter-FLuc-CMV-RLuc-dsRed2), the firefly luciferase-CMV promoter-renilla luciferase-dsRed2 sequence was subcloned from the pDR5-fluc-CMV-RlucDsRed2 vector (gift from Dr Khalid Shah; (21)) using primer pairs targeting the relevant promoter sequence (**Table 1**), which contained a SalI and XhoI site. The DNA fragment underwent restriction enzyme digest (New England Biolabs) to generate overhangs and was subcloned into the pCCL vector. Specific promoter regions upstream of the transcription start site (**Table 1**) were then subcloned into the editable promoter-reporter construct. The DNA fragments were PCR-amplified with primer pairs containing a XhoI and BamHI site from human genomic DNA (Promega) and subcloned into the pCLL-NoPromoter-FLuc-CMV-RLuc-dsRed2 vector. Complete insertion of the promoter sequence was confirmed by Sanger sequencing (GENEWIZ) and whole plasmid sequencing (Full Circle Labs).

**Table 1:**
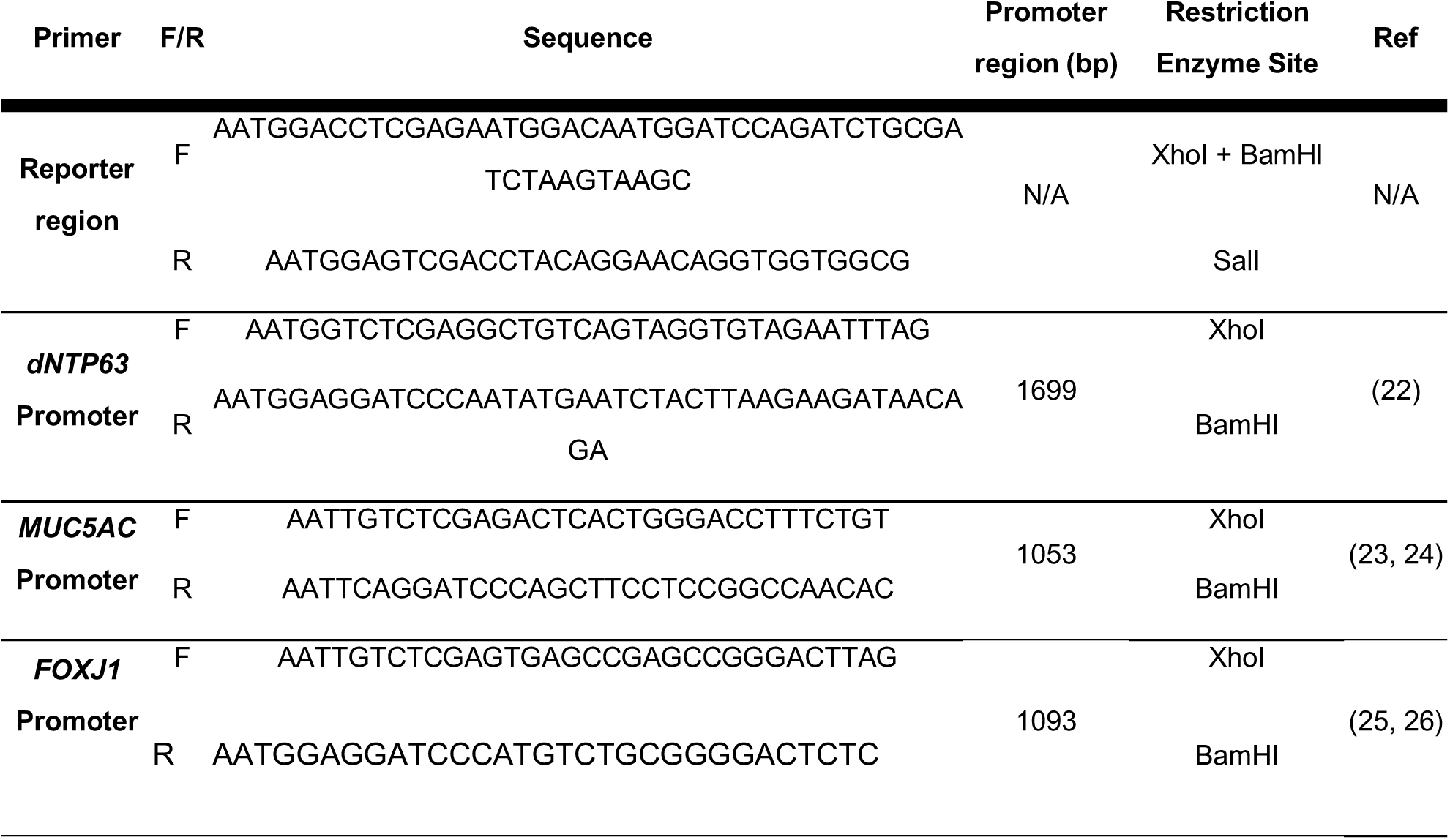
Oligonucleotide primers used for construct generation.

### Data visualization

Schematics were created in Inkscape (v1.2.1). Analyses and data visualization were performed in RStudio (v2023.12.0.369) with the tidyverse (v2.0.0; (27)) packages: dplyr (v1.1.4), tidyr (v1.3.0), tibble (v3.2.1) ggplot2 (v3.4.4), ggpubr (v.0.6.0) and the colorblind-friendly color map (Viridis, v0.6.4; (28)) .

Genomic location and sequences of genes were visualized using the Gviz package (v1.46.1; (29)). Gene information was extracted from the ENSEMBL hg38 genome using biomaRt (v2.58.0; (30, 31)) and filtered for the HUGO gene nomenclature committee symbol.

### Cell lines

HEK293Ts were cultured in DMEM with pyruvate (Gibco) supplemented with 10% FBS and 1 x penicillin/streptomycin for one passage prior to and during transfection. The HBEC3-KT cell line (ATCC) was cultured in Airway Epithelial Cell Basal Medium (ATCC) supplemented with the Bronchial Epithelial Cell Growth Kit (ATCC) or in Keratinocyte SFM (Gibco), a serum-free medium supplemented with recombinant EGF and bovine pituitary extract. The OE19 cell line (a kind gift from The Francis Crick Institute Cell Services STP) and the CAPAN-2 cell line (a gift from Prof Hemant Kocher, Barts Cancer Institute, London, U.K.) were cultured in RPMI (Gibco) supplemented with 10% FBS (Gibco) and 1 x penicillin/streptomycin (Gibco). Cells were passaged using 0.05% trypsin-EDTA. Cells were centrifuged at 300 x g for five min to ensure trypsin removal.

### Lentivirus production

Viral supernatants were created by co-transfecting HEK293T cells at 70-80% confluency in a T175 with 20 µg lentiviral construct, 13 µg pCMVR8.74 (gift from Didier Trono, Addgene plasmid #22036; http://n2t.net/addgene:22036; RRID:Addgene_22036) and 7 µg pMD2.G (gift from Didier Trono, Addgene plasmid #12259; http://n2t.net/addgene:12259; RRID:Addgene_12259) using JetPEI (Polyplus Transfection) following the manufacturer’s protocol. Viral supernatants were collected 48 and 72 hours post-transfection and filtered through a 0.45 µm filter (Whatman). Supernatant was combined with PEGit concentrator (5X; System Biosciences) overnight at 4°C and centrifuged at 1500 g for 45 min at 4°C. The supernatant was removed, and the viral pellet was resuspended in 1/10^th^ of the original supernatant volume in DMEM with pyruvate supplemented with 25 mM HEPES (Gibco). Concentrated supernatants were stored at -80°C until use.

### Primary human airway epithelial cell culture

3T3-J2 mouse embryonic fibroblasts (Kerafast) were expanded in DMEM with pyruvate containing 9% bovine serum and 1X penicillin/streptomycin (Gibco). These cells were mitotically inactivated by treatment with 4 µg/mL mitomycin C (Sigma-Aldrich) for three hours to produce feeder layers. Cells were trypsinized and plated at a density of 20,000 cells/cm^2^. Epithelial cells were added the following day in primary epithelial cell culture medium, as previously described (32, 33). Primary epithelial cell culture medium consisted of DMEM with pyruvate (Gibco) and F12 (Gibco) in a 3:1 ratio with 1X penicillin/streptomycin and 5% FBS supplemented with 5 μM Y-27632 (Cambridge Bioscience), 25 ng/mL hydrocortisone (Sigma), 0.125 ng/mL EGF (Sino Biological), 5 μg/mL insulin (Sigma), 0.1 nM cholera toxin (Sigma), 250 ng/mL amphotericin B (Fisher Scientific) and 10 μg/mL gentamicin (Gibco). The differential trypsin sensitivity of 3T3-J2 cells and epithelial cells allows the removal of feeder cells on passage. A second period of trypsinization allows passage of the epithelial cells.

### Lentiviral transduction

The HBEC3-KT cell line, CAPAN-2 cell line and primary airway epithelial cells were transduced upon passaging. 100 μL of concentrated virus were added to 150,000 cells in suspension in a total volume of 500 μL. Cells were agitated every 5 min for 30 min to keep the cells in suspension and then plated for expansion in appropriate medium. Medium was replaced after overnight adherence of the cells. The OE19 cell line was plated at 150,000 cells per well in a 6-well plate, the following day these cells were transduced with the addition of 100 μL concentrated virus and polybrene (4 μg/mL; Santa Cruz) to the cell medium. Medium was replaced 7 hours after the addition of the virus.

### Fluorescence-activated cell sorting (FACS)

Transduced cells were enriched by FACS to purify the dsRed2^+^ populations. Single cell suspensions of cells were generated by trypsinization. Differential trypsinization was performed for primary cell cultures to remove 3T3-J2 feeder cells. Cells were filtered through a 70 µm strainer (Falcon), centrifuged at 300 x g for 5 min, and resuspended in FACS buffer consisting of 1% FBS, 25 mM HEPES and 1 mM EDTA in PBS. Sorting was performed on either a BD FACS Aria or a BD FACS Aria Fusion.

### Dual luciferase assay

The firefly and renilla luciferase activities were confirmed in the transduced cell populations using a dual luciferase assay. The transduced cell lines and untransduced controls (HBEC3-KT, OE19 and CAPAN-2 cells) were seeded at 10,000 cells per well in a 384-well plate (Thermo Scientific) and after 24 h the luciferase activities were determined using the Dual-Glo Luciferase Assay System (Promega) according to the manufacturer’s protocol. The relative light unit was measured on an Envision II plate reader.

### Air-liquid interface culture

Air-liquid interface (ALI) culture followed a previously published method (34). Briefly, 0.4 μm PET transwells (Falcon) in 24-well plates (Falcon) were coated for one hour with 50 μg/mL Collagen I (Corning) in 0.02N acetic acid. Transwells were washed twice with PBS and air dried for 15 min. Epithelial cells were seeded at 300,000 cells/transwell in 250 μL of primary airway epithelial cell culture medium. 700 μL of primary airway epithelial cell culture medium was added to the basolateral side of the transwell. On day 2, medium was aspirated from the apical side of the transwell, exposing the confluent layer of epithelial cells to air. Medium was replaced on the basolateral side with PneumaCult medium (STEMCELL Technologies) which was prepared as per the manufacturer’s instructions. Medium was replaced twice a week and the apical transwell side was washed once a week with PBS to remove accumulated mucus. ALI cultures underwent bioluminescence imaging on days 2, 9, 16, 23 and 30 and were collected for RNA isolation and fixed for immunofluorescence staining. For RNA isolation, membranes were washed twice with PBS and cells were lysed on the membrane using RLT plus buffer and processed following the manufacturer’s protocol with the addition of an extra elution step to maximize RNA concentration (RNeasy Plus Mini Kit; Qiagen). RNA was stored at -80 °C prior to use.

### Quantitative real-time polymerase chain reaction

A NanoDrop One^C^ (Thermo Scientific) was used to quantify RNA concentration. 500 ng RNA was converted to cDNA using qScript cDNA SuperMix (Quanta Bio) following the manufacturer’s protocol in a 20 µL reaction. cDNA samples were diluted 1 in 2 and stored at - 20 °C prior to use. Quantitative real-time polymerase chain reaction (qPCR) was performed on a QuantStudio™ 5 Real-Time PCR System (Applied Biosystems) with the Power SYBR Green PCR Master Mix (Applied Biosystems). Oligonucleotide primers (**Table 2**) were used at 200 nM and technical triplicates were tested. Relative RNA quantification was calculated by the delta Ct method with glyceraldehyde 3-phosphate dehydrogenase (*GAPDH*) and ribosomal protein S13 (*RPS13*) as reference genes (**Table 2**). Fold change was calculated by the delta-delta Ct method.

**Table 2:**
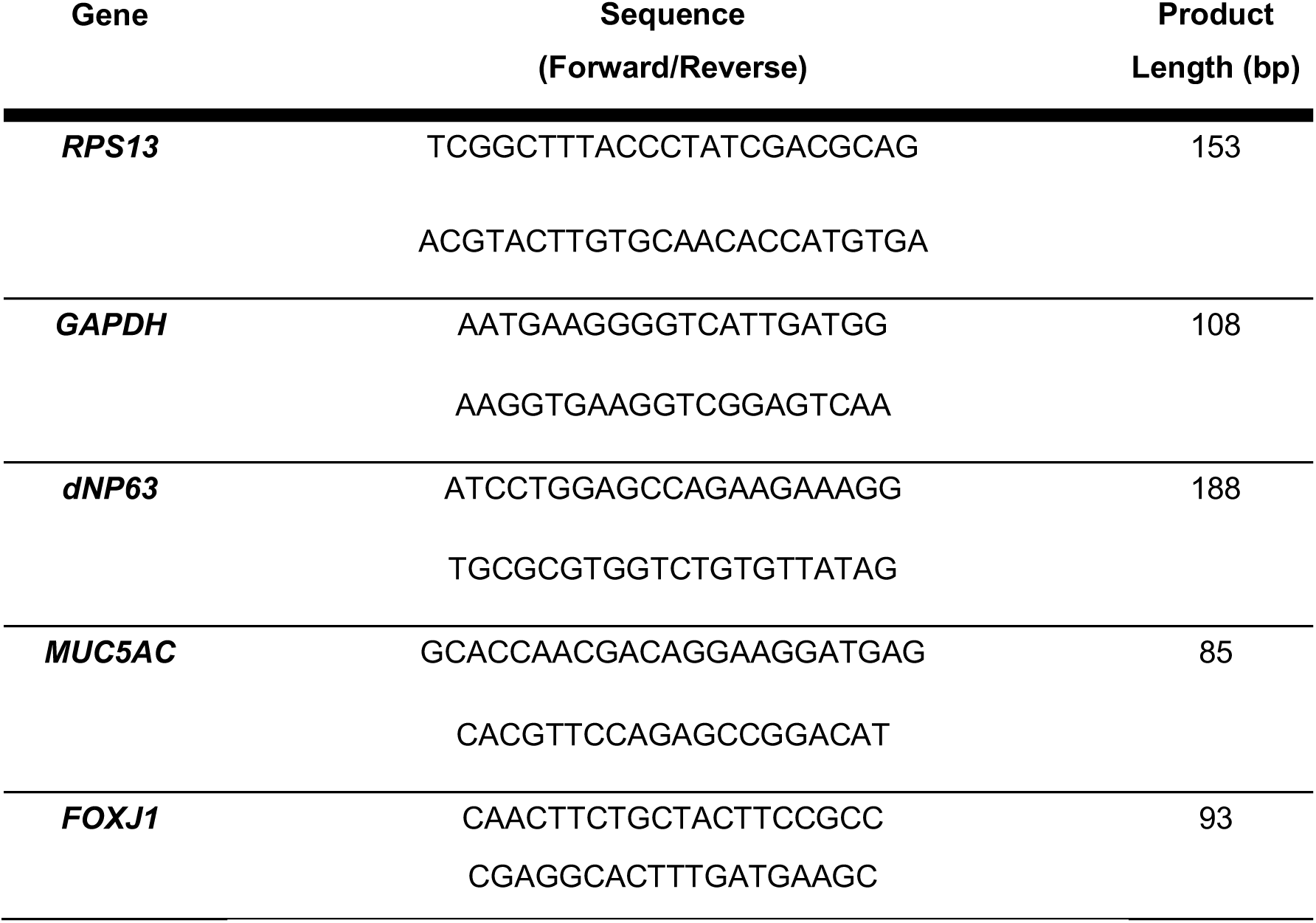
Oligonucleotide primers for qPCR

### 3D organoid (‘bronchosphere’) culture

Bronchosphere culture followed a previously published method (32). Differentiation medium consisted of 50 % DMEM with pyruvate and 50 % BEBM supplemented with BEGM singlequots (except amphotericin B, triiodothyronine, and retinoic acid) (Lonza). Medium was supplemented with 100 nM all-trans retinoic acid (Sigma-Aldrich) at time of use. Briefly, wells of an ultra-low attachment 96-well plate (Corning) were coated with 30 μL of 25% matrigel (Corning) in differentiation medium and returned to the incubator at 37°C for 30 min. Epithelial cells were seeded at 2,500 cells/well in 65 μL of 5% matrigel in differentiation medium containing 5 μM Y-27632.

Bronchospheres were fed with 50 μL differentiation medium on days 3, 10 and 17 of culture. Where bronchospheres were supplemented with cytokines, human recombinant interleukin-6 (IL-6; PeproTech) or human recombinant IL-13 (PeproTech) were included in medium additions on days 3, 10 and 17, and control wells received medium containing an equivalent amount of BSA. Bronchospheres were collected on day 24 in ice cold PBS and centrifuged at 300 x g for 5 min, then fixed with 4% PFA on ice for 30 min and centrifuged at 400 x g for 5 min.

Bronchospheres were washed with ice cold PBS and transferred to a well of a V-bottomed 96-well plate (Thermo Scientific). The plate was centrifuged at 400 x g for 5 min, the supernatant was removed, and Bronchospheres were resuspended in HistoGel (Epredia). After 10 min on ice, the gels were transferred to 70% ethanol before tissue processing using standard procedures.

### Bioluminescence imaging

Bioluminescence imaging was conducted on an IVIS Spectrum (PerkinElmer) within the Centre for Advanced Biomedical Imaging at UCL. D-luciferin (Abcam) was added to cell cultures at 150 µg/mL with 25 mM HEPES in DMEM with pyruvate. For ALI cultures, differentiation medium was removed from the basolateral side of the transwell and replaced with DMEM with pyruvate containing luciferin and HEPES; 200 µL was added to the apical side of the transwell. For bronchosphere cultures, 10 µL luciferin-containing medium was spiked into the well containing differentiation medium, giving the same final concentration as in the ALI cultures, as medium removal would disturb the 3D structures.

Bioluminescent and brightfield images were taken at regular intervals from 10 min after luciferin addition until after the peak of the bioluminescent signal was recorded. This was typically within 30 min. For ALI cultures, the luminescence imaging was taken with a 1 min exposure time, medium binning and an aperture (f/stop) of 1. For bronchosphere cultures, the bioluminescence imaging was taken with a 2 min exposure time, medium binning, and an aperture (f/stop) of 1. For all cultures, brightfield images were taken with medium binning and an aperture (f/stop) of 8.

### Histology and immunofluorescence staining

Samples were embedded in paraffin type 6 wax (Epredia) using an embedding station (Sakura Tissue-TEK TEC) and 5 µm sections were cut on a Microm HM 325 microtome. Haematoxylin and eosin (H&E) staining was performed on sections using an automated staining system (Sakura Tissue-Tek DRS) and imaged on a Nanozoomer Digital Pathology (Hamamatsu). Histology images were collated and presented from whole-slide image files using PATHOverview (available on GitHub (https://github.com/EpiCENTR-Lab/PATHOverview).

For immunofluorescence staining of samples on slides, slides were dewaxed using an automated protocol, washed in PBS and a hydrophobic ring was drawn around the sample using an ImmEdge pen (Vector Laboratories). Sections were blocked with 1% bovine serum albumin (BSA; Merck), 5 % normal goat serum (NGS; Abcam) and 0.1 % Triton X-100 (Sigma-Aldrich) in PBS for 1 hour at room temperature. Primary antibodies (**Table 3**) were diluted in block buffer and applied to slides overnight at 4 °C. Slides were washed twice with PBS. Secondary antibodies conjugated to species-appropriate AlexaFluor dyes were diluted 1:1000 in 5 % NGS, 0.1 % Triton X-100 in PBS and applied to slides for 3 hours at room temperature in the dark. 100 ng/mL DAPI (Sigma-Aldrich) in PBS was applied to the slides for 20 min. Slides were washed twice in PBS and a coverslip was applied manually with Immu-Mount (Thermo Scientific). Immunofluorescence images were acquired using a Leica DMi8 microscope. To automate quantification analyses, macros were created in Fiji (35) to mask and quantify the nuclei count and area of immunofluorescent staining.

**Table 3:** Primary antibodies used for immunofluorescence staining.

| Antibody | Catalog number (Supplier) | Clone | Dilution factor |
| --- | --- | --- | --- |
| TP63 | Ab124762 (Abcam) | Rabbit | 1:300 |
| KRT5 | 905901 (Biolegend) | Chicken | 1:500 |
| MUC5AC | M5293 (Sigma) | Mouse IgG1 | 1:500 |
| ACT | T6793 (Sigma) | Mouse IgG2b | 1:500 |
| FOXJ1 | 14-9965-82 (ThermoFisher Scientific) | Mouse IgG1 | 1:200 |

## RESULTS

### pCLL-NoPromoter-FLuc-CMV-RLuc-dsRed2: a versatile lentiviral luciferase gene reporter construct

To generate a lentiviral reporter system suitable for monitoring airway cell type-specific gene expression, we first generated a construct in which we could insert gene-specific promoter sequences (using the XhoI and BamHI sites) to drive firefly luciferase expression and have renilla luciferase and the dsRed2 fluorescent protein expressed from the constitutively active CMV promoter (pCLL-NoPromoter-FLuc-CMV-RLuc-dsRed2; **Figure 1A (left)**, henceforth “No Promoter”).

**Figure 1:**
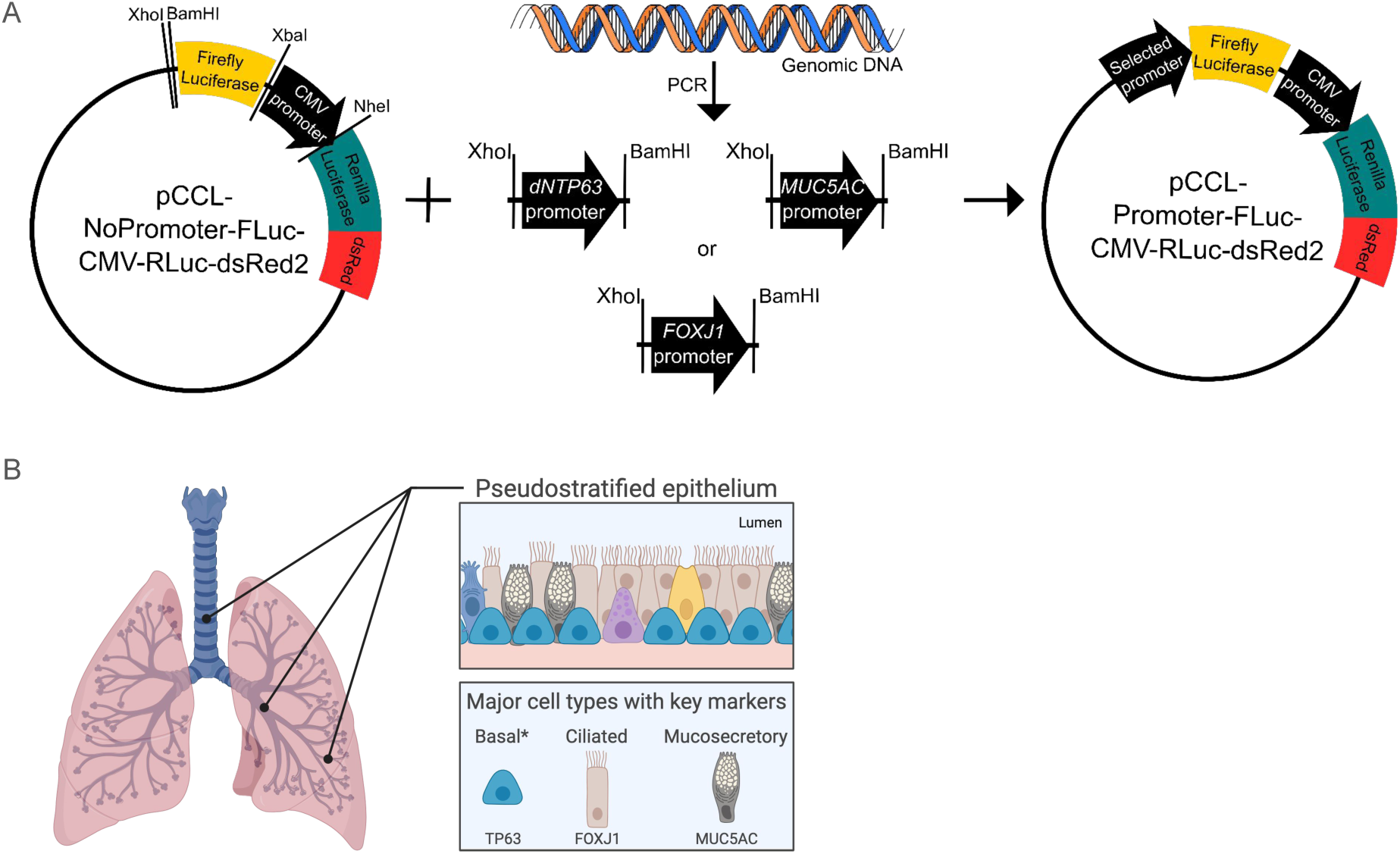
Development of a lentiviral gene reporter system for bioluminescence monitoring of airway cell type-specific gene expression. **A)** The pCCL-NoPromoter-FLuc-CMV-RLuc-dsRed2 lentiviral construct has no promoter sequence upstream of the firefly luciferase gene. Renilla luciferase and dsRed2 are under the control of the constitutively active CMV promoter. XhoI and BamHI restriction sites allow insertion of selected promoter sequences upstream of firefly luciferase. **B)** Schematic representation of the human airway epithelium. In this study, we inserted promoter sequences from the *dNTP63*, *MUC5AC* and *FOXJ1* genes into independent lentiviral constructs to create reporters for basal, mucosecretory and ciliated cell differentiation, respectively. Panel B was created using BioRender.com.

Next, we aimed to use this vector to produce cell type-specific promoter-reporter constructs for the main cell types of the pseudostratified airway epithelium (**Figure 1B**); basal, mucosecretory and ciliated cells. We identified sequences with literature evidence of promoter activity that were upstream of the transcription start sites of the *TP63*, *MUC5AC* and *FOXJ1* genes, respectively (22–26). Each promoter sequence was isolated by PCR of genomic DNA with primer pairs that included the XhoI and BamHI sequences. This enabled subcloning of each sequence into the No Promoter construct (**Figure 1A**). Promoter sequence insertion was first confirmed by restriction enzyme digestion of the constructs using the XhoI and BamHI enzymes and validated by Sanger sequencing and whole plasmid sequencing (sequences and plasmid maps are available via Addgene).

### A *dNTP63* promoter-reporter construct to monitor basal cell differentiation

The TP63 gene has multiple isoforms (**Figure 2A**), but airway basal cells mainly express ΔNp63α. As such the *dNTP63* promoter sequence was used to generate the pCLL-dN*TP63*Promoter-FLuc-CMV-RLuc-dsRed2 (henceforth, “*dNTP63* reporter”) construct (**Figure 2A**). To ensure that this construct faithfully reports the expression of *TP63*, we first transduced the HBEC3-KT cell line with lentivirus carrying the dN*TP63* reporter vector or the No Promoter control reporter vector. After FACS to purify transduced cells based on dsRed2 expression, expression of both firefly and renilla luciferases were detected in the cells transduced with the *dNTP63* reporter and only renilla luciferase was detected in cells transduced with the No Promoter control vector (**Figure 2B**).

**Figure 2:**
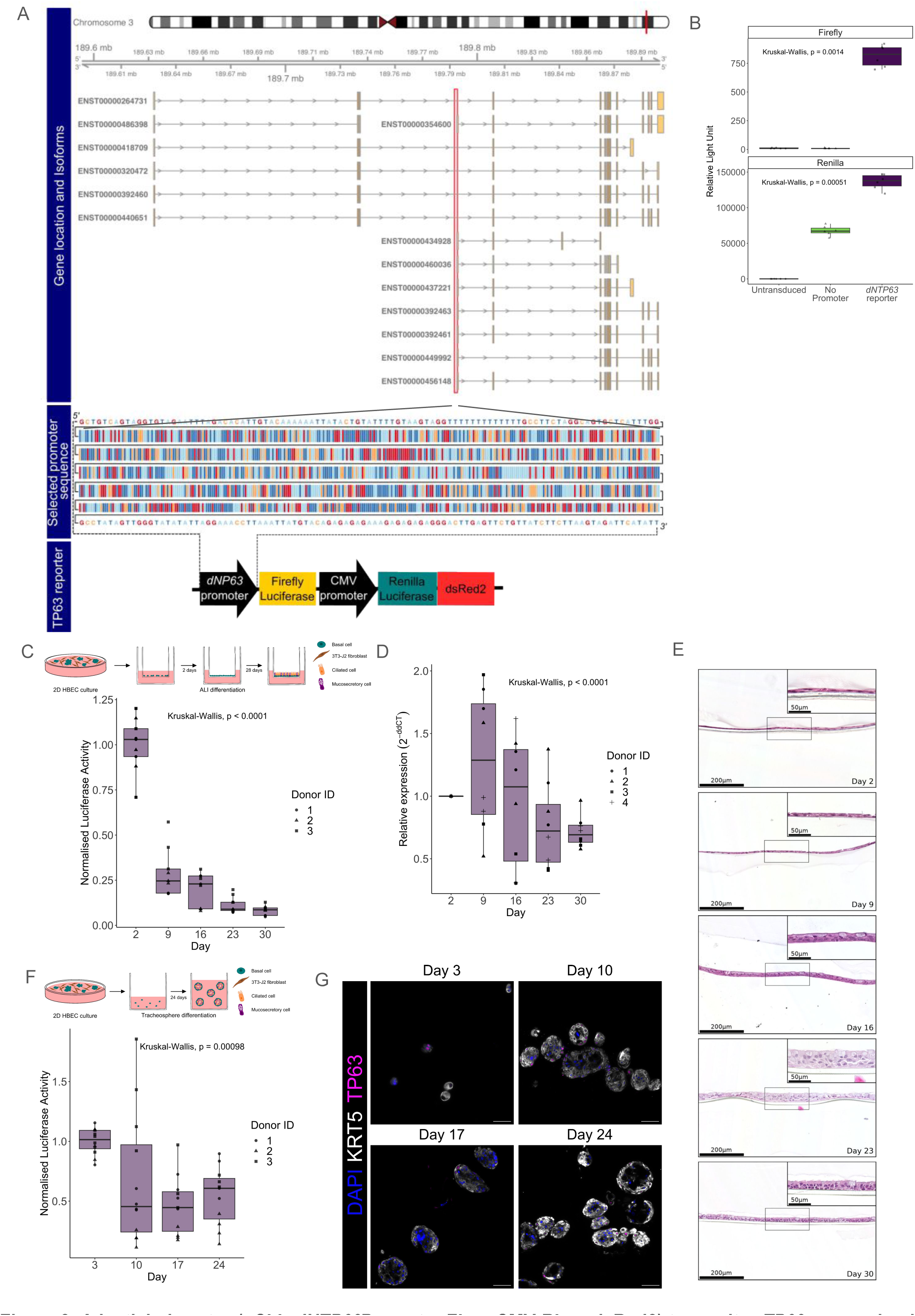
A lentiviral vector (pCLL-*dNTP63*Promoter-FLuc-CMV-RLuc-dsRed2) to monitor *TP63* expression in human airway epithelial cells. **A)** Design of *dNTP63* promoter reporter vector. Genomic location of *TP63* and *TP63* isoforms (upper panel). Selected *dNTP63* promoter sequence is highlighted in a red box and shown (middle panel). The final *dNTP63* reporter construct (pCLL-dN*TP63*Promoter-FLuc-CMV-RLuc-dsRed2) is shown (lower panel). **B)** Quantitation of firefly and renilla bioluminescence (relative light units) in HBEC3-KT cells transduced with either the *dNTP63* reporter or No Promoter reporter. Data shown with transduced cells from a single transduction, each data point is a single well reading. **C)** Quantification of *dNTP63* promoter driven firefly luciferase bioluminescence over time in the air liquid interface (ALI) model using primary human bronchial epithelial cells (HBECs) transduced with the *dNTP63* reporter (n=9, 3 primary cell donors with triplicate wells per donor across 2 independent experiments). Data normalized to the mean peak average radiance (p/s/cm^2^/sr) on day 2 of the differentiation assay per donor. **D)** qPCR quantification of *dNTP63* transcript expression over time in all promoter reporter transduced primary HBEC ALI cultures (n=11/12, 4 primary cell donors; each data point is the mean of technical triplicates from a single transduction with promoter-reporter constructs). **E)** Representative haematoxylin and eosin stained images of ALI sections over time from dNTP63 reporter transduced primary human bronchial epithelial cells (Donor ID 3). Figure created with PATHOverview. **F)** Quantification of *dNTP63* promoter driven firefly luciferase bioluminescence over time in the bronchosphere differentiation assay using primary HBECs transduced with the *dNTP63* reporter (n=9, 3 primary cell donors with triplicate wells per donor across 2 independent experiments). Data normalized to the mean peak average radiance (p/s/cm^2^/sr) on day 3 of the differentiation assay for each donor. **G)** Representative immunofluorescence images of bronchosphere sections over time from a single donor of *dNTP63* reporter transduced primary human bronchial epithelial cells (Donor ID 3) of basal cell markers keratin 5 (KRT5) and TP63. Scale bars = 50 μm. Statistical analyses were performed with the Kruskal-Wallis test.

Next, we transduced primary human bronchial epithelial cells with lentivirus carrying the dN*TP63* reporter vector. Transduction efficiency ranged from 5.2-41.0% (mean = 22.5%, n = 6 donors). After FACS to purify transduced cells based on dsRed2 expression, we seeded the transduced basal cells in air-liquid interface cultures and monitored luciferase expression during a 30 day time course after culture initiation (**Figure 2C**). We observed that dN*TP63* reporter activity declined over time (**Figure 2C**), as expected based on the differentiation of TP63^+^ basal cells to TP63^-^ luminal cell types in this assay (36). Following the luciferase assay, wells were collected for end-point analyses including qPCR which confirmed this result (**Figure 2D**).

Further, these air-liquid interface cultures were collected for histological staining to validate and visualize differentiation of transduced airway basal cells to a pseudostratified airway epithelium (**Figure 2E**). We also examined the behavior of the reporter during differentiation in 3D bronchosphere cultures (**Figure 2F**). Detection of firefly luciferase decreased as differentiation of bronchospheres proceeded (**Figure 2F**), and immunofluorescence staining for basal cell markers confirmed that the proportion of TP63^+^ cells decreased over time (**Figure 2G**).

### A *MUC5AC* promoter-reporter construct to monitor mucosecretory cell differentiation

To validate that the pCCL-*MUC5AC*Promoter-FLuc-CMV-RLuc-dsRed2 (henceforth, “*MUC5AC* reporter”) construct (**Figure 3A**), we transduced OE-19 cells, an esophageal adenocarcinoma cell line that was identified as expressing *MUC5AC* in the Human Protein Atlas cell line database (https://www.proteinatlas.org/). Activity of both firefly and renilla luciferase activity was detected in the transduced cell line (**Figure 3B**), so we proceeded to transduce primary human bronchial epithelial cells. Transduction efficiency ranged from 3.3-33.5% (mean = 11.0%, n = 6 donors). Following FACS purification of transduced cells, we observed a decrease in luciferase expression in multiple post-airlift air-liquid interface culture timepoints (**Figure 3C**). This was despite seeing the expected increase in *MUC5AC* expression in qPCR experiments (**Figure 3D**) and increased MUC5AC-positivity in immunofluorescence staining of sections from air-liquid interface cultures (**Figure 3E**). In 3D bronchospheres derived from transduced basal cells, *MUC5AC* reporter expression also did not change (p = 0.841, two-way ANOVA). However, when we introduced the cytokine IL-13 to bronchosphere cultures to increase mucosecretory cell differentiation and expression of MUC5AC (37, 38), the luciferase signal was significantly increased (p (Condition) = 0.0000012, two-way ANOVA) (**Figure 3F**). Assessment of MUC5AC protein levels in bronchospheres by immunofluorescence revealed low abundance during standard differentiation but much higher abundance in IL-13-treated cultures at day 17 (**Figures 3G and 3H**), potentially explaining the lack of induction of the firefly luciferase reporter in standard differentiation conditions.

**Figure 3:**
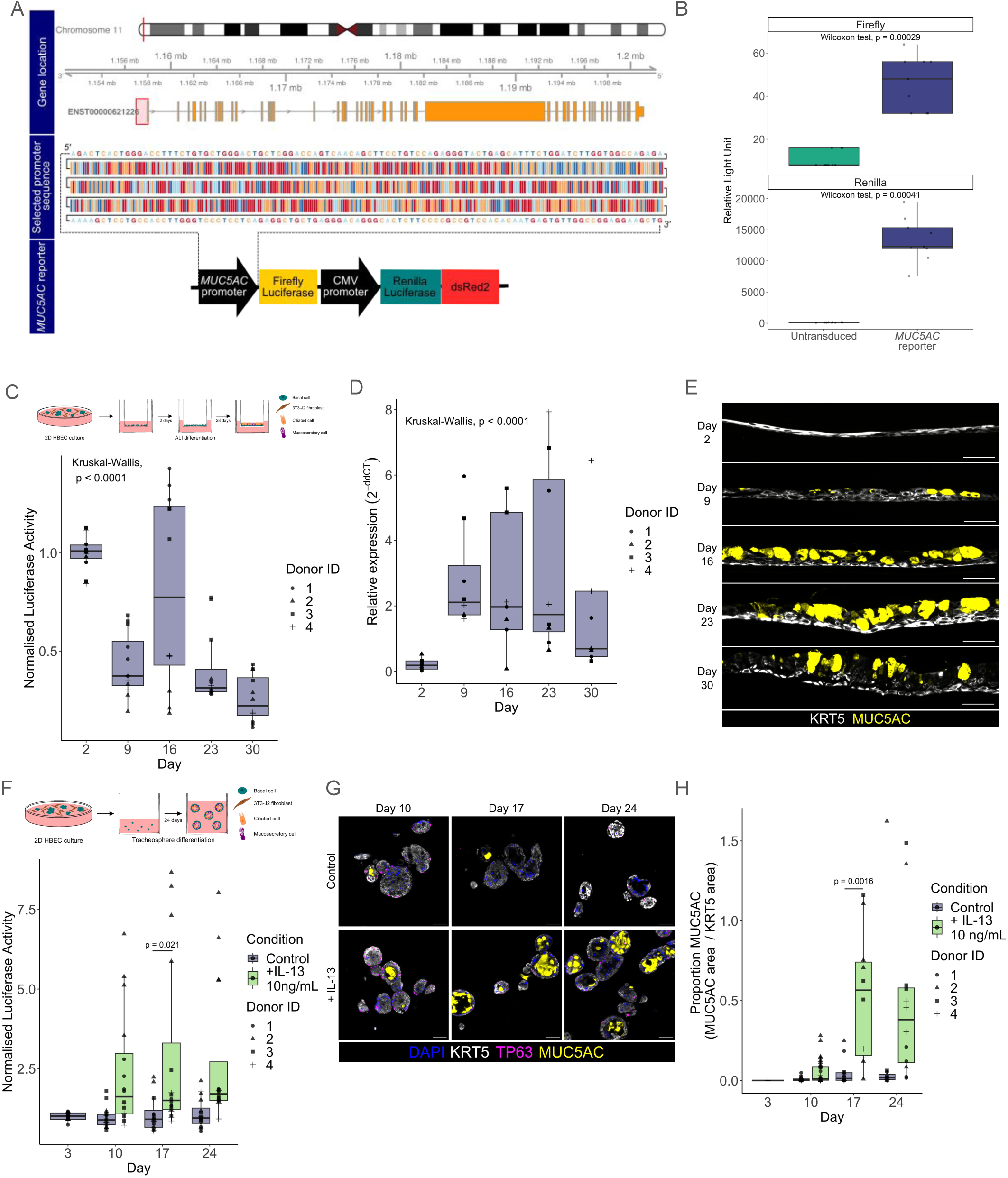
A lentiviral vector (pCCL-*MUC5AC*Promoter-FLuc-CMV-RLuc-dsRed2) to monitor *MUC5AC* expression in human airway epithelial cells. **A)** Design of *MUC5AC* promoter reporter vector. Genomic location of *MUC5AC* (upper panel). Selected *MUC5AC* promoter sequence is highlighted in a red box and shown (middle panel). The final *MUC5AC* reporter construct (pCLL-*MUC5AC*Promoter-FLuc-CMV-RLuc-dsRed2) is shown (lower panel). **B)** Quantitation of firefly and renilla bioluminescence (relative light units) in OE-19 cells transduced with the *MUC5AC* reporter. Data shown with transduced cells from a single transduction, each data point is a single well reading. A Wilcoxon non-parametric test was performed. **C)** Quantification of *MUC5AC* promoter driven firefly luciferase bioluminescence over time in the air liquid interface (ALI) model using primary human bronchial epithelial cells (HBECs) transduced with the *MUC5AC* reporter (n=12, 4 primary cell donors with triplicate wells per donor across 2 independent experiments). Data normalized to the mean peak average radiance (p/s/cm^2^/sr) on day 2 of the differentiation assay for each donor. The Kruskal-Wallis test was performed. **D)** qPCR quantification of *MUC5AC* transcript expression over time in all promoter reporter transduced primary HBEC ALI cultures (n=6-8, 4 primary cell donors; each data point is the mean of technical triplicates from a single transduction with promoter-reporter constructs. A Kruskal-Wallis test was performed. **E)** Immunofluorescence staining of ALI cultures of HBECs transduced with the *MUC5AC* promoter reporter (keratin 5 (KRT5), white; MUC5AC, yellow; representative section images of time course cultures from donor 3. Scale bars = 50 μm. **F)** Quantification of *MUC5AC* promoter driven firefly luciferase bioluminescence over time in bronchospheres cultured in the presence of 10 ng/mL IL-13 or a BSA control using primary HBECs transduced with the *MUC5AC* reporter (n=16, 4 primary cell donors; each data point is a single well reading across 2 independent experiments). Data normalized to the mean peak average radiance (p/s/cm^2^/sr) on day 3 of the differentiation assay for each donor. A two-way ANOVA was performed, p (overall) = 0.841, p (condition) = 0.0000012, p (day) = 0.191. Significant Tukey test p-values are reported between conditions on each day. **G)** Immunofluorescence staining of bronchosphere cultures of *MUC5AC* reporter transduced HBECs cultured in the presence of 10 ng/mL IL-13 or a BSA control (keratin 5 (KRT5), white; TP63, magenta; MUC5AC, yellow; representative section images of time course cultures from donor 3. Scale bars = 50 μm. **H)** Quantification of the immunofluorescence staining of bronchosphere cultures of *MUC5AC* reporter transduced HBECs cultured in the presence of 10 ng/mL IL-13 or a BSA control. n=4-29, 4 HBEC donors, each data point is from a single image of bronchosphere sections. A two-way ANOVA was performed, p (overall) = 0.006279, p (condition) = 0.000225, p (day) = 0.0000733. Significant Tukey test p-values are reported between conditions on each day.

### A *FOXJ1* promoter-reporter construct to monitor multiciliated cell differentiation

To validate that the pCLL-*FOXJ1*Promoter-FLuc-CMV-RLuc-dsRed2 (henceforth, “*FOXJ1* reporter”) construct (**Figure 4A**), we transduced CAPAN-2 cells, a pancreatic ductal adenocarcinoma cell line that was identified as expressing *FOXJ1* in the Human Protein Atlas cell line database (https://www.proteinatlas.org/). Firefly luciferase activity was detected following transduction and FACS was performed based on dsRed2 expression (**Figure 4B**). In primary cell cultures, we observed lentiviral transduction efficiencies of 6.1-31.0% (mean = 12.8%, n = 6 donors). Following FACS purification of transduced cells, we observed increased *FOXJ1* reporter luciferase expression at days 16, 23 and 30 compared to days 2 and 9 of air-liquid interface culture (**Figure 4C**), consistent with the emergence of FOXJ1^+^ multiciliated cells in air-liquid interface cultures after day 12 post-airlift in these cultures (11). We validated increased *FOXJ1* transcript abundance in our cultures using qPCR (**Figure 4D**). In 3D bronchospheres derived from transduced basal cell cultures, we also saw increased expression of FOXJ1 during the differentiation period (p = 0.0142, two-way ANOVA) (**Figure 4E**). Addition of IL-6 to bronchosphere cultures, which is known to promote ciliated cell differentiation *in vitro* (39), led to increased firefly luciferase signal at days 17 and 24 of culture, timepoints at which ciliated cells are expected to be present in cultures (**Figure 4E**). Quantification of immunofluorescence staining for FOXJ1 confirmed that ciliated cell numbers increased during bronchosphere differentiation, and that IL-6 increased FOXJ1^+^ ciliated cell abundance (**Figure 4F, 4G**).

**Figure 4:**
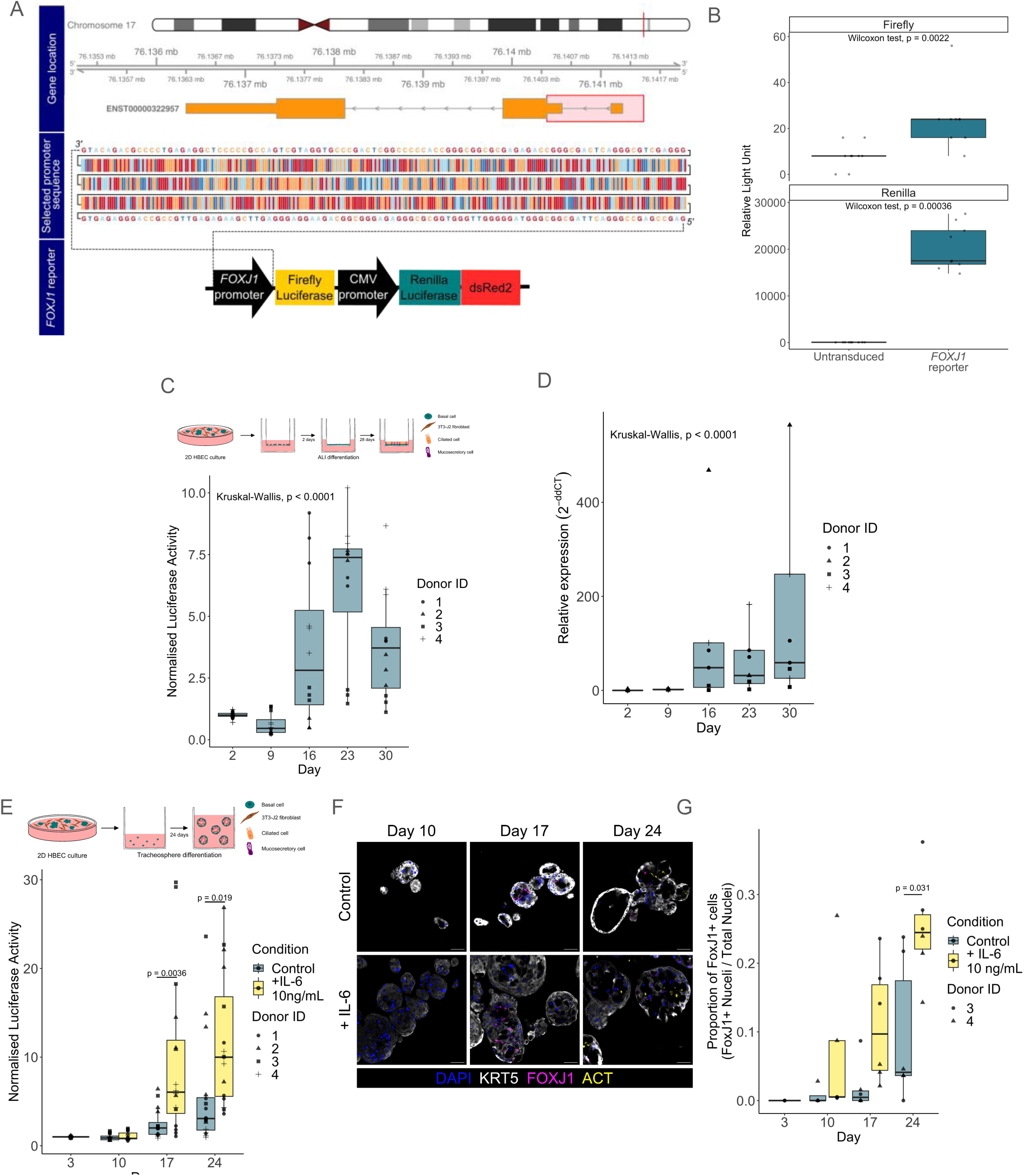
A lentiviral vector (pCLL-FOXJ1Promoter-FLuc-CMV-RLuc-dsRed2) to monitor *FOXJ1* expression in human airway epithelial cells. **A)** Design of *FOXJ1* promoter reporter vector. Genomic location of *FOXJ1* (upper panel). Selected *FOXJ1* promoter sequence is highlighted in a red box and shown (middle panel). The final *FOXJ1* reporter construct (pCLL-*FOXJ1*Promoter-FLuc-CMV-RLuc-dsRed2) is shown (lower panel). **B)** Quantitation of firefly and renilla bioluminescence (relative light units) in CAPAN-2 cells transduced with the *FOXJ1* reporter. Data shown with transduced cells from a single transduction, each data point is a single well reading. A Wilcoxon non-parametric test was performed. **C)** Quantification of *FOXJ1* promoter driven firefly luciferase bioluminescence over time in the air liquid interface (ALI) model using primary human bronchial epithelial cells (HBECs) transduced with the *FOXJ1* reporter (n=12, 4 primary cell donors with triplicate wells per donor across 2 independent experiments). Data normalized to the mean peak average radiance (p/s/cm^2^/sr) on day 2 of the differentiation assay for each donor. **D)** qPCR quantification of *FOXJ1* transcript expression over time in all promoter reporter transduced primary HBEC ALI cultures (n=7/8, 4 primary cell donors; each data point is the mean of technical triplicates from a single transduction with promoter-reporter constructs. **E)** Quantification of *FOXJ1* promoter driven firefly luciferase bioluminescence over time in bronchospheres cultured in the presence of 10 ng/mL IL-6 or a BSA control using primary HBECs transduced with the *FOXJ1* reporter (n=16, 4 primary cell donors; each data point is a single well reading across 2 independent experiments). Data normalized to the mean peak average radiance (p/s/cm^2^/sr) on day 3 of the differentiation assay for each donor. A two-way ANOVA was performed, p (overall) = 0.0142, p (condition) = 0.0000396, p (day) = 0.0000000154. Significant Tukey test p-values are reported between conditions on each day. **F)** Immunofluorescence staining of bronchosphere cultures of *FOXJ1* reporter transduced HBECs cultured in the presence of 10 ng/mL IL-6 or a BSA control (keratin 5 (KRT5), white; FOXJ1, magenta; ACT, yellow; representative section images of time course cultures from donor 3. Scale bars = 50 μm. **G)** Quantification of the immunofluorescence staining of bronchosphere cultures of *FOXJ1* reporter transduced HBECs cultured in the presence of 10 ng/mL IL-6 or a BSA control. n=4-6, 4 HBEC donors, each data point is from a single image of bronchosphere sections. A two-way ANOVA was performed, p (overall) = 0.415, p (condition) = 0.000367, p (day) = 0.000525. Significant Tukey test p-values are reported between conditions on each day.

## DISCUSSION

To overcome some of the limitations of existing approaches for studying airway epithelial cell differentiation, and to enable real-time monitoring of the differentiation process, we developed lentiviral gene reporter constructs for basal cells, mucosecretory cells and multiciliated cells. We developed a construct in which a cell type-specific promoter sequence drives expression of firefly luciferase, and the CMV promoter drives expression of both renilla luciferase and dsRed2, a red fluorescent protein. dsRed2 allows facile FACS isolation of a pure population of transduced cells, while firefly luciferase can be normalized to renilla luciferase to achieve a per cell quantification of promoter activity.

We validate that promoter sequences from *TP63*, *MUC5AC* and *FOXJ1* correlated with the abundance of relevant cell populations in primary air-liquid interface (ALI) and 3D bronchosphere cultures. *dNTP63* promoter activity declines in ALI cultures, reflecting the reduction in expression of *dNTP63* transcripts per cell during the emergence of *TP63*^-^ differentiated epithelial cells. Further, in the bronchosphere assay the decline in *dNTP63* promoter activity over time is consistent with the transition from single HBECs to 3D structures containing both TP63^+^KRT5^+^ basal cells at the periphery and TP63^-^ differentiated cells on the luminal surface.

We observed a relative reduction in *MUC5AC* promoter activity in ALI cultures from day two to day nine which was in contrast to the qPCR data and the emergence of MUC5AC protein over time. There is evidence that submerged HBECs upregulate several secretory and mucin genes, including *MUC5AC*, in our culture conditions compared to culture in BEGM (33). This may explain the high luciferase activity recorded at day two of the differentiation assay. Despite the unexpected trajectory of luciferase activity in the ALI differentiation of *MUC5AC* reporter HBECs, the addition of interleukin 13 (IL-13) to the bronchosphere assay, a cytokine which is known to induce mucosecretory differentiation (37, 38), resulted in an increase in luciferase activity on day 17 and 24. This mirrored the increase of MUC5AC observed in bronchospheres cultured with IL13, confirming that this lentiviral construct could be used in applications to identify modulators of airway mucosecretory differentiation.

The expected increase in *FOXJ1* promoter activity as the differentiation assays progressed was observed by bioluminescence imaging. These results reflected the increase in *FOXJ1* transcripts in ALI culture and the emergence of FOXJ1^+^ nuclei in the bronchosphere assay. The increase of ciliated cell abundance by addition of IL-6 (39) to the bronchosphere cultures was observed by an increase in luciferase activity on day 17 and 24, validating the use of this lentiviral construct to identify modulators of airway ciliated cell differentiation.

The non-destructive nature of the luciferase assay on live cell cultures enables repeated measurements to be made from the same cultures over a time course, with these cultures then available for additional end-point assays (such as TEER readings, ciliary beat frequency analyses, qPCR and immunofluorescence). Further, the lentiviral constructs reported here could be useful tools in iPSC-derived airway epithelial cell differentiation protocols to provide real-time monitoring of cell type emergence and/or to act as a quality control measure between cultures (40, 41). Given the existence of multiple luciferases that luminesce at different wavelengths (42), in the future it may be possible to generate a lentiviral vector that monitors multiple airway epithelial cell populations simultaneously. Moreover, the cloning strategy used here created a versatile vector (pCCL-NoPromoter-FLuc-CMV-RLuc-dsRed2) into which a promoter sequence (or candidate promoter sequence) from any gene of interest can readily be inserted.

The lentiviruses developed here have potential applications in compound screening in primary cell cultures from healthy and disease patient populations. Since *TP63* expression correlates with stemness in stratified squamous epithelia (43), the reporter might be used to identify modulators of epithelial stemness, for example. The *FOXJ1* reporter could see application in investigations aiming to identify modulators of multiciliated cell differentiation (44), while the *MUC5AC* reporter is relevant in various disease contexts. Goblet cell metaplasia or hyperplasia is characterized by increased differentiation of MUC5AC^+^ mucosecretory cells. Since cigarette smoke exposure induces MUC5AC expression in air-liquid interface cultures (45), screening compounds for their ability to reduce smoking-induced mucosecretory cell differentiation is feasible using this approach. Similarly, airway epithelial cells from patients with asthma and COPD have higher expression levels of MUC5AC than those from healthy individuals (46–48), so screens to identify compounds to reduce goblet cell metaplasia or mucous hypersecretion are now also feasible in disease contexts.

## DATA AVAILABILITY STATEMENT

All relevant data are made available with this manuscript. Plasmids for the lentiviral constructs described are available via Addgene: pCCL-NoPromoter-FLuc-CMV-RLuc-dsRed2 (Addgene #215329), pCCL-dNTP63Promoter-FLuc-CMV-RLuc-dsRed2 (Addgene #215326), pCCL-MUC5ACPromoter-FLuc-CMV-RLuc-dsRed2 (Addgene #215327) and pCCL-FOXJ1Promoter-FLuc-CMV-RLuc-dsRed2 (Addgene #215328).

## ACKNOWLEDGEMENTS

The authors thank Mr Jamie Evans (Flow Cytometry Core Facility; UCL Division of Medicine, University College London, London, U.K.) and Mr George Morrow (UCL Cancer Institute Flow Cytometry Facility, University College London, London, U.K.) for their assistance with FACS experiments. The authors also thank Bernadette Carroll (Department of Thoracic Medicine, University College Hospital, London, U.K.), Dr Matthew Wright (Department of Thoracic Medicine, University College Hospital, London, U.K.) and members of the ASCENT study team (UCL Respiratory, University College London, U.K.) for their roles in collecting patient tissue used in this study. Finally, the authors thank members of the Epithelial Cell Biology in ENT Research (EpiCENTR) group (University College London, U.K.) for feedback on a draft manuscript.

## DISCLAIMERS

The views expressed are those of the authors and not necessarily those of the NHS, the NIHR or the Department of Health. For the purpose of open access, the author has applied a CC BY public copyright license to any author accepted manuscript version arising from this submission.

## AUTHOR CONTRIBUTIONS

J.C.O. and R.E.H. conceived and designed research; J.C.O, A.L., P.F.D., M.-B.E.M. and K.A.L. performed experiments; J.C.O. analyzed data; J.C.O., S.M.J. and R.E.H. interpreted results of experiments; J.C.O. prepared figures: J.C.O. and R.E.H. drafted the manuscript; J.C.O., M.-B.E.M., S.M.J. and R.E.H. edited and revised the manuscript: S.M.J. and R.E.H. approved the final version of the manuscript.

